# A NanoBRET-based binding assay for Smoothened allows real time analysis of small-molecule ligand binding and distinction of two separate ligand binding sites for BODIPY-cyclopamine

**DOI:** 10.1101/706028

**Authors:** Paweł Kozielewicz, Carl-Fredrik Bowin, Gunnar Schulte

## Abstract

**Background and Purpose:** Smoothened (SMO) is a GPCR that mediates hedgehog signaling. Hedgehog binds the Patched, which in turn regulates SMO activation. Overactive SMO signaling is oncogenic and is therefore a clinically established drug target. Here, we establish a nanoluciferase bioluminescence resonance energy transfer (NanoBRET)-based ligand binding assay for SMO providing a sensitive and high throughput-compatible addition to the toolbox of GPCR pharmacologists.

**Experimental Approach:** In the NanoBRET-based binding assay, SMO is N terminally tagged with nanoluciferase (Nluc) and binding of BODIPY-cyclopamine is assessed by quantifying resonance energy transfer between receptor and ligand. The assay allows kinetic analysis of ligand-receptor binding in living HEK293 cells and competition binding experiments using commercially available SMO ligands (SANT-1, cyclopamine-KAAD, SAG1.3 and purmorphamine).

**Key Results:** The NanoBRET binding assay for SMO is sensitive and superior to purely fluorescence-based binding assays. BODIPY-cyclopamine showed complex binding parameters suggesting separate binding sites.

**Conclusions and Implications:** The NanoBRET ligand binding assay for SMO provides a fast, sensitive and reliable alternative to assess SMO ligand binding. Furthermore, this assay is sufficiently sensitive to dissect a SANT-1-sensitive and a SANT-1-insensitive cyclopamine binding site in the 7TM core, and will be important to further dissect and understand the molecular pharmacology of Class F receptors.

**What is already known:** Cyclopamine targets SMO as antagonist and fluorescently-labelled cyclopamine has been used for fluorescence-based binding assays for SMO. Structural analysis has suggested two binding sites on SMO, one in the receptor core and one the CRD.

**What this study adds:** We established a NanoBRET-based binding assay for SMO with superior sensitivity compared to fluorescence-based assays. This assay allows distinction of two separate binding sites for BODIPY-cyclopamine on SMO in live cells in real time.

**What is the clinical significance:** The assay is a valuable complement for drug discovery efforts and will support a better understanding of Class F GPCR pharmacology.

## 1. INTRODUCTION

Smoothened (SMO) is a G protein-coupled receptor (GPCR) which, alongside ten paralogues of Frizzleds (FZDs), forms the Class F of GPCRs (Schulte, 2010). SMO signaling is of utmost importance during embryonic patterning and development, and dysfunction of SMO signaling is causative in the development of diverse tumors including basal cell carcinoma (Ingham & McMahon, 2001). Therefore, pharmacological targeting of SMO and SMO signaling evolved as an attractive antitumor treatment strategy established in clinical practice (Wu, Zhang, Sun, McMahon & Wang, 2017). On a structural level, this seven transmembrane domain spanning receptor is characterized by a large, extracellular cysteine-rich domain (CRD) and a long C-terminal domain (Schulte, 2010). While SMO is essential for transmitting transcriptional responses via heterotrimeric G proteins and Glioma-associated oncogene (Gli) signaling induced by hedgehog proteins (three mammalian homologues: Desert, Indian and Sonic hedgehog), the nature and mode of action of the endogenous ligand and the mechanisms of receptor activation are not fully understood (Byrne et al., 2016; Schulte & Kozielewicz, 2019). It is known that hedgehog proteins bind Patched, a cholesterol transporter, which in turn regulates SMO activation by postulated regulation of cholesterol levels (Zhang et al., 2018). Cholesterol and other naturally occurring sterols are positive allosteric modulators and agonists of SMO in Gli- and G protein-dependent signaling (Nachtergaele et al., 2012; Qi, Liu, Thompson, McDonald, Zhang & Li, 2019; Raleigh et al., 2018; Sever et al., 2016). Due to the distinct link to human cancer and occurrence of several cancer-associated SMO mutations (e.g. D473H^6.54^ or W535L^7.55^; superscript numbering refers to Ballesteros Weinstein nomenclature of GPCRs (Ballesteros & Weinstein, 1995)), a plethora of small ligands – antagonists and inverse agonists – have been developed to target this receptor (Wu, Zhang, Sun, McMahon & Wang, 2017). Two of these compounds – vismodegib and sonidegib - are approved as drugs for the treatment of basal-cell carcinoma (Chen, 2016). In addition, nature provides an effective SMO antagonist, the plant alkaloid cyclopamine (Incardona, Gaffield, Kapur & Roelink, 1998; Taipale et al., 2000).

These and other ligands, such as SMO agonists (e. g. SAG1.3, purmorphamine), neutral antagonists (e. g. cyclopamine, SANT-1) or inverse agonists (e. g. cyclopamine-KAAD), are frequently used to explore SMO pharmacology (Chen, 2016; Chen, Taipale, Young, Maiti & Beachy, 2002; Rominger et al., 2009). To date, binding affinities of these compounds were determined using classical radioligand binding methods (Frank-Kamenetsky et al., 2002; Rominger et al., 2009; Wang et al., 2014) and, more often, fluorescence-based assays using for example the green-yellow fluorescent BODIPY-cyclopamine (Chen, Taipale, Cooper & Beachy, 2002; Chen, Taipale, Young, Maiti & Beachy, 2002; Gorojankina et al., 2013; Huang et al., 2016; Huang et al., 2018; Manetti et al., 2010). The fluorescently-labelled cyclopamine has been used in three separate experimental approaches: assessment of ligand-receptor interaction based on detection of fluorescence using confocal microscopy, flow cytometry or fluorescence polarization (Bee et al., 2012; Chen, Taipale, Cooper & Beachy, 2002; Chen, Taipale, Young, Maiti & Beachy, 2002; Gorojankina et al., 2013; Huang et al., 2016; Huang et al., 2018). While these methods offer valuable insight into ligand-receptor binding in live cells, they suffer from several shortcomings, including: 1) laborious protocols that require long ligand incubation times (up to 4-5 hours), 2) extensive cell washing due to lipophilicity of BODIPY-cyclopamine, 3) high levels of non-specific binding in untransfected cells or in the presence of saturating concentrations of unlabeled competitors and 4) the need for data normalization of BODIPY-fluorescence values to receptor expression values.

In order to overcome these experimental limitations, we established and validated a live-cell, nanoluciferase bioluminescence resonance energy transfer (NanoBRET)-based binding assay to assess the binding properties of BODIPY-cyclopamine and unlabeled SMO ligands to an N-terminally Nanoluciferase (Nluc)-tagged SMO and ΔCRD SMO in HEK293 cells devoid of endogenous SMO. This proximity-based ligand-binding assay has recently been developed to assess ligand binding to Class A GPCRs and receptor tyrosine kinases (Bosma et al., 2019; Bouzo-Lorenzo et al., 2019; Mocking, Verweij, Vischer & Leurs, 2018; Stoddart et al., 2015; Stoddart, Kilpatrick & Hill, 2018; Stoddart et al., 2018; Sykes, Stoddart, Kilpatrick & Hill, 2019). It relies on the high specificity of BRET between Nluc-tagged protein (BRET donor) and fluorescently-tagged ligand (BRET acceptor) that can only occur when both BRET partners are within a distance of 10 nm (100 Å). Thus, the interference of non-specifically bound probe, outside of the BRET radius to the receptor is – in contrast to detecting solely ligand fluorescence – minimal. In the present study, we employ NanoBRET to monitor binding of commercially available SMO ligands. Furthermore, the sensitivity of the assay enables us to dissect the pharmacological properties of separate cyclopamine binding sites in the transmembrane-spanning receptor core of the Class F receptor SMO. Thus, this NanoBRET-based binding assay provides a valuable complement to the toolbox of high-throughput compatible screening assays for Class F GPCRs.

## 2. METHODS

### 2.1 Cell culture

ΔSMO HEK293 cells were generated and characterized as described elsewhere (Kozielewicz et al., 2019, submitted). The cells were cultured in DMEM (HyClone) supplemented with 10% FBS, 1% penicillin/streptomycin, and 1% L-glutamine (all Thermo Fisher Scientific) in a humidified CO2 incubator at 37 °C. All cell culture plastics were from Sarstedt, unless otherwise specified.

#### DNA cloning and mutagenesis

Nluc-A3 was from Stephen Hill, University of Nottingham, UK (Stoddart et al., 2015). SMO-*R*luc8 coding for mouse Smoothened was from Nevin A. Lambert, Augusta University, Georgia, Canada (Wright et al., 2019). The mouse SMO sequence was subcloned into an empty N-terminally tagged Nluc vector containing 5-HT_3_A signal peptide using BamHI and XbaI restriction sites. First, the BamHI site present in mouse SMO was removed using site-directed mutagenesis (GeneArt, Thermo Fisher Scientific) with the following primers: 5’-CCTCCAGGGGCTGGGGTCCATTCATTCCCGC-3’ (forward primer) and 5’-GCGGGAATGAATGGACCCCAGCCCCTGGAGG-3’ (reverse primer). Next, the mouse SMO sequence was cloned in-frame into the Nluc vector using forward primer: 5’-GAC GGA TCC GCG GCC TTG AGC GGG AAC GTG-3’ and reverse primer: 5’-CGT TCT AGA TCA GAA GTC CGA GTC TGC ATC-3’ . ΔCRD Nluc-SMO was generated using the mouse SMO lacking the BamHI site by cloning it into N-terminally tagged Nluc vector between BamHI and XbaI using forward primer: 5’-GAC GGA TCC GAG GTA CAA AAC ATC AAG TTC-3; and reverse primer: 5’-CGT TCT AGA TCA GAA GTC CGA GTC TGC ATC-3’ . ΔCRD and full-length Nluc-SMO D477G^6.54^/E522K^7.38^ were generated with site-directed mutagenesis (GeneArt, Thermo Fisher Scientific) by first obtaining D477G^6.54^ mutation with the following primers: 5’-GCTGCCACTTCTATGGCTTCTTCAACCAGGC-3’ (forward primer) and 5’-GCCTGGTTGAAGAAGCCATAGAAGTGGCAGC-3’ (reverse primer). Subsequently the E522K^7.38^ mutation was introduced with 5’-CCCAGCCTCCTGGTGAAGAAGATCAATCTAT-3’ (forward primer) and 5’-ATAGATTGATCTTCTTCACCAGGAGGCTGGG-3 (reverse primer). All constructs were validated by sequencing (Eurofins GATC, Konstanz, Germany).

### 2.2 Live-cell ELISA

For quantification of cell surface receptor expression by labelling with anti-Nluc antibody, ΔSMO HEK293 cells at the density of 4·10^5^ cells ml^−1^ were transfected in suspension using Lipofectamine 2000 with 50-500 ng of the indicated receptor plasmid DNA with 500-950 ng of pcDNA plasmid DNA. The cells (100 μl) were seeded onto a PDL-coated transparent 96-well plate with flat bottom and grown overnight. 24 hr later the cells were washed twice with 0.5% BSA in PBS and incubated with a rabbit anti-Nluc (2 μg·ml^−1^; RnD Systems #MAB10026) in 1% BSA/PBS for 1 hr at 4°C. Following incubation, the cells were washed four times with 0.5% BSA/PBS and incubated with a horseradish peroxidase-conjugated goat anti-rabbit antibody (1:3,000; Thermo Fisher Scientific #31460) in 1% BSA/PBS for 1 hr at 4°C. The cells were washed three times with 0.5% BSA/PBS, and 50 μl of the peroxidase substrate TMB (3,3’,5,5’-tetramethylbenzidine; Sigma-Aldrich #T8665) were added. The cells were further incubated for 20 minutes and upon development of a blue product, 50 μl of 2 M HCl were added and the absorbance was read at 450 nm using a BMG Ω POLARstar plate reader. The data were analyzed in GraphPad Prism 6.

### 2.3 Immunoblotting

ΔSMO HEK293 cells were transfected in suspension using Lipofectamine 2000 (50-500 of receptor plasmid DNA with 500-950 ng of pcDNA plasmid DNA per 4·10^5^ cells ml^−1^) and seeded (700 μl) onto wells of a 24-well plate. Protein lysates were obtained using Laemmli buffer with 0.5% NP-40 and 5% β-mercaptoethanol. Lysates were sonicated and analyzed on 4–20 % Mini-PROTEAN TGX precast polyacrylamide gels (Bio-Rad) and transferred to PVDF membranes using the Trans-Blot Turbo system (Bio-Rad). After blocking with 5% milk in TBS-T, membranes were incubated with primary antibodies in blocking buffer: rabbit anti-GAPDH (1:5000; Cell Signaling Technology #2118) and mouse anti-Nluc (0.5 μg·ml^−1^; RnD Systems #MAB10026), overnight at 4 °C. Proteins were detected with horseradish peroxidase-conjugated secondary antibody (1:5,000; goat anti-rabbit (Thermo Fisher Scientific #31460 or 1:3,000; goat anti-mouse; Thermo Fisher Scientific #31430) and Clarity Western ECL Blotting Substrate (Bio-Rad).

### 2.4 NanoBRET binding assay

ΔSMO HEK293 cells were transiently transfected in suspension using Lipofectamine 2000 (Thermo Fisher Scientific). 4·10^5^ cells ml^−1^ were transfected with 50-500 ng of receptor plasmid DNA and 500-950 ng of pcDNA. The cells (100 μl) were seeded onto a PDL-coated black 96-well cell culture plate with solid flat bottom (Greiner Bio-One). 24 hr post-transfection, cells were washed once with HBSS (HyClone) and maintained in the same buffer. In the saturation experiments, the cells were incubated with different concentrations of BODIPY-cyclopamine (80 μl) for 60 min at 37°C before the addition of the luciferase substrate coelenterazine h (5 μM final concentration, 10 μl) for 6 min prior to the BRET measurement. In the competition experiments, the cells were pre-incubated with different concentrations of unlabeled ligands (70 μl) for 30 min at 37°C. Fixed concentration of BODIPY-cyclopamine was then added (10 μl) and the cells were incubated for additional 60 min at 37°C before the addition of the luciferase substrate colenterazine h (5 μM final concentration, 10 μl) for 6 min prior to the BRET measurement. In the association experiments, the cells were pre-incubated with 10 μM SANT-1 (30 minutes), followed by colenterazine h (5 μM final concentration) at 37°C prior to the addition of different BODIPY-cyclopamine concentrations. The BRET signal was measured every minute for 90 min at 37°C. The BRET ratio was determined as the ratio of light emitted by BODIPY-cyclopamine (energy acceptor) and light emitted by Nluc-tagged biosensors (energy donors). The BRET acceptor (bandpass filter 535–30 nm) and BRET donor (bandpass filter 475–30 nm) emission signals were measured using a CLARIOstar microplate reader (BMG). ΔBRET ratio was calculated as the difference in BRET ratio of cells treated with ligands and cells treated with vehicle. BODIPY fluorescence was measured prior to reading luminescence (excitation: 477–14 nm, emission: 525–30 nm). Data were analyzed using GraphPad Prism 6.

### 2.5 Data and statistical analysis

Data were analyzed using GraphPad Prism 6 and represent mean ± standard error of the mean (SEM) or standard deviation (SD) of n independent experiments (biological replicates) performed typically in triplicates (technical replicates). Please refer to the figure legends for more details on the displayed data.

BODIPY-cyclopamine saturation curves were fitted using three-parameter or biphasic nonlinear regression models (logarithmic scale for BODIPY-cyclopamine concentrations) or one- or two-site saturation nonlinear regression models (linear scale for BODIPY-cyclopamine concentrations). Models were selected based on an extra-sum-of square F-test (p<0.05).

Competition binding curves were analyzed using a one-site binding model in order to obtain equilibrium dissociation constants values (K_i_) of unlabeled ligands as per the Cheng-Prusoff equation (Cheng & Prusoff, 1973):

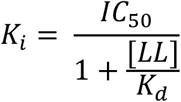

In the equation: K_i_ is the searched dissociation constant of an unlabeled ligand, IC_50_ is the inhibitory constant 50 of an unlabeled ligand obtained from the competition curve, [LL] is the concentration of a labelled ligand used in the competition experiment and K_d_ is the equilibrium dissociation constant of a labelled ligand obtained from the saturation studies.

To analyze the labelled ligand binding kinetics data, one-phase association or two-phase association models were selected based on an extra-sum-of square F-test:

One-phase association:

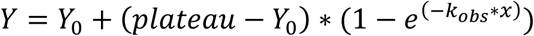

Two-phase association:

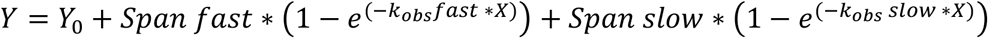

Where: Y_0_ is Y value at time x = 0, plateau is the Y value at infinite times and k_obs_ is the association constant expressed in 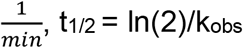

Given the observed complexity of BODIPY-cyclopamine binding at different concentrations, herein we present the raw values from kinetics experiments for each concentrations at the tested SMO constructs.

### 2.6 Materials

BODIPY-cyclopamine was from BioVision Inc. Purmorphamine (9-Cyclohexyl-N-[4-(4-morpholinyl)phenyl]-2-(1-naphthalenyloxy)-9H-purin-6-amine) was from Abcam. SAG1.3 (3-Chloro-N-[trans-4-(methylamino)cyclohexyl]-N-[[3-(4-pyridinyl)phenyl]methyl]benzo[b]thiophene-2-carboxamide) was from Sigma. Cyclopamine-KAAD and SANT-1 ((4-Benzyl-piperazin-1-yl)-(3,5-dimethyl-1-phenyl-1H-pyrazol-4-ylmethylene)-amine) were from Abcam. All ligands were dissolved in DMSO and stored in aliquots at −20°C. The ligands underwent a maximum of 2 freeze-thaw cycles. Colenterazine h was from Biosynth and it was stored as 2.4 mM aliquots in acidified ethanol at −80°C. Protein-low binding tubes (Eppendorf) were used to make serial dilutions of BODIPY-cyclopamine.

## 3. RESULTS

### N-terminally Nluc tagged SMO constructs are expressed at the cell surface

In order to establish a NanoBRET-based binding assay for the Class F receptor SMO, we adopted the cloning strategy of previously presented Class A GPCR including a 5-HT_3_A receptor-derived signal sequence and an extracellular, N-terminally Nluc fused to either the full length mouse SMO or SMO lacking the extracellular cysteine-rich domain (CRD). Subsequently, these constructs are referred to as Nluc-SMO and ΔCRD Nluc-SMO, respectively (**Fig. 1a**). Both receptor constructs are expressed in the cells and at the cell surface upon transient transfection of ΔSMO HEK293 cells as shown by immunoblotting of whole cell lysates and a live-cell surface ELISA (**Fig. 1a, b**).

**Fig. 1:**
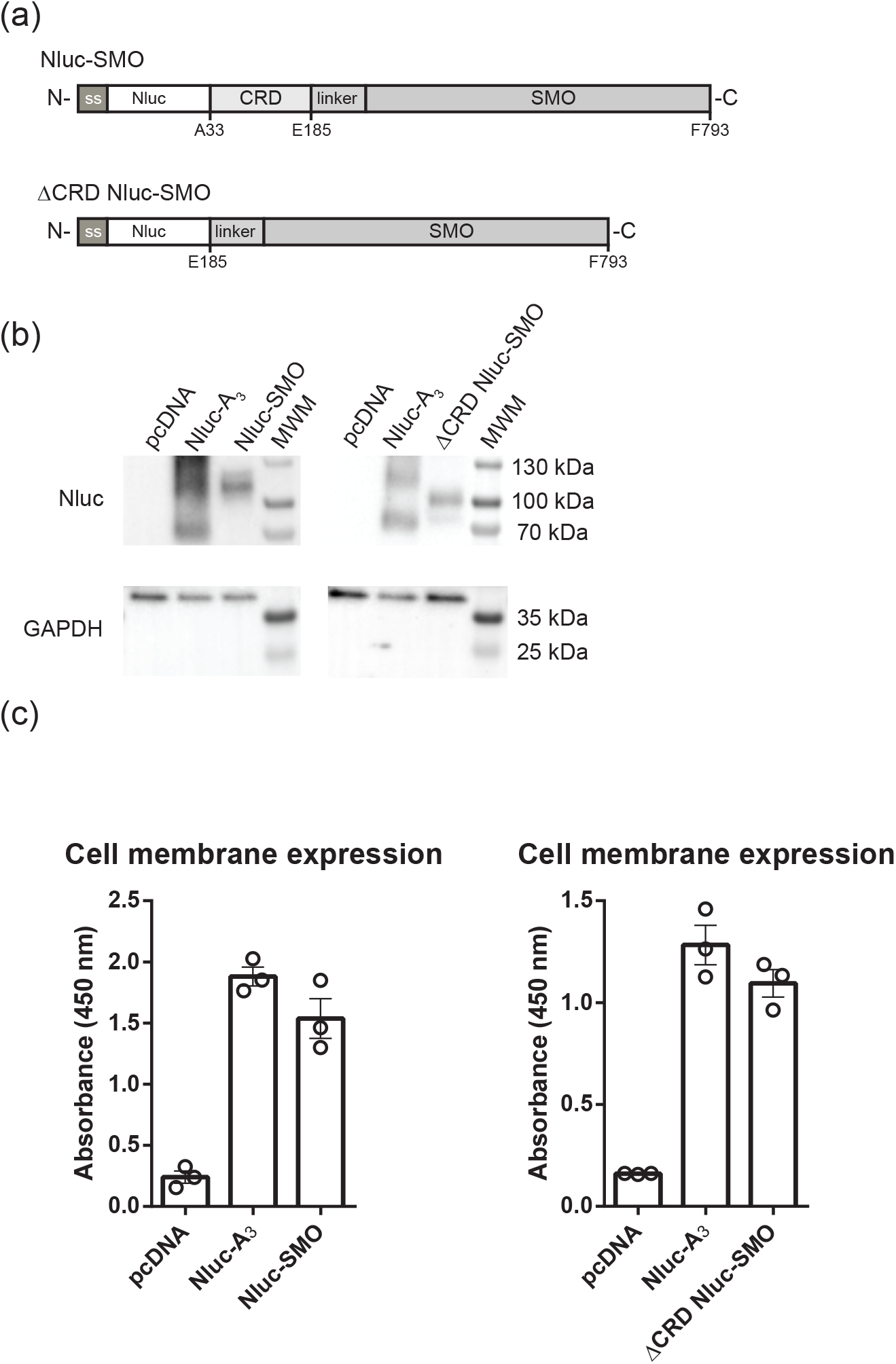
Construct validation of the N terminally tagged Nluc-SMO. (a) Schematic presentation of the NanoBRET Nluc-SMO and ΔCRD Nluc-SMO sensors for BODIPY-cyclopamine binding. (b) Validation of cellular expression of Nluc-SMO and ΔCRD Nluc-SMO upon transient transfection into ΔSMO HEK293 cells. Cell lysates were analyzed by immunoblotting using anti-Nluc antibody and anti-GAPDH served as a loading control. The higher apparent molecular weights of Nluc-A3 (positive control, predicted molecular weight = 55 kDa), Nluc-SMO (predicted molecular weight = 103 kDa) and ΔCRD Nluc-SMO (predicted molecular weight = 87 kDa) could be a result of N-glycosylation of the receptors. The experiments were repeated three times with similar results. (c) Surface expression of Nluc-SMO and ΔCRD Nluc-SMO was quantified by ELISA based on labelling with an anti-Nluc antibody. Raw data are shown, N=3.

### BODIPY-cyclopamine binding to Nluc-SMO can be monitored by NanoBRET

The commercially available, BODIPY-labelled derivative of the plant alkaloid cyclopamine (BODIPY-cyclopamine; **Fig. 2a**) associates with Nluc-SMO transiently expressed in ΔSMO HEK293 cells in a concentration-dependent manner reaching saturation at ~1000 nM (one-site fit K_d_ = 170.9 ± 26.6 nM, **Fig. 2b, c**). Similarly, BODIPY-cyclopamine binds the SMO construct lacking the CRD (two-sites fit K_d1_ = 4.0 ± 0.9 nM, **Fig. 2d,e**). The BODIPY-cyclopamine bound ΔCRD Nluc-SMO with higher affinity and importantly the NanoBRET produced by BODIPY-cyclopamine binding was larger in the ΔCRD Nluc-SMO compared to Nluc-SMO (two-sites fit ΔBRET B_max1_ ΔCRD receptor = 0.08 ± 0.01 and one-site fit ΔBRET B_max_ full-length receptor = 0.034 ± 0.001). Furthermore, BODIPY-cyclopamine binding to ΔCRD Nluc-SMO was more complex than binding to Nluc-SMO as indicated by the multiphasic binding curve for ΔCRD Nluc-SMO especially at higher concentrations of the ligand (two-sites fit K_d2_ = 371.3 ± 266.1 nM) . While quantification of BODIPY-cyclopamine binding on living cells was assessed 60 min after ligand addition, we were also interested in the binding kinetics of BODIPY-cyclopamine. Therefore, we followed BODIPY-cyclopamine association at different ligand concentrations using Nluc-SMO and ΔCRD Nluc-SMO expressing ΔSMO HEK293 cells (**Fig. 3a,b**). First, the kinetic analysis confirmed that saturation was reached at 60 min ligand exposure as used for experiments presented in Fig. 2. In agreement with our observations from saturation experiments, the kinetic binding analysis underlined that the affinity of BODIPY-cyclopamine to the ΔCRD Nluc-SMO was higher and BRET counts were larger compared to Nluc-SMO. Reflecting the different parameters of the BODIPY-cyclopamine binding to the two different SMO constructs, we used 100, 200, 1000 nM ligand for Nluc-SMO binding kinetics and 10, 100, 1000 nM for ΔCRD Nluc-SMO. Moreover, the analysis also showed a substantial difference in t_1/2_ of BODIPY-cyclopamine at both receptors (**Table 1** and **2**). In order to further dissect BODIPY-cyclopamine association to SMO we employed the SMO antagonist SANT-1 at 10 μM, which interacts with the 7TM core of the receptor (Chen, Taipale, Young, Maiti & Beachy, 2002). For both receptor constructs SANT-1 reduced association of BODIPY-cyclopamine at the two lower concentrations but did not completely abrogate binding of 1000 nM BODIPY-cyclopamine.

**Fig. 2:**
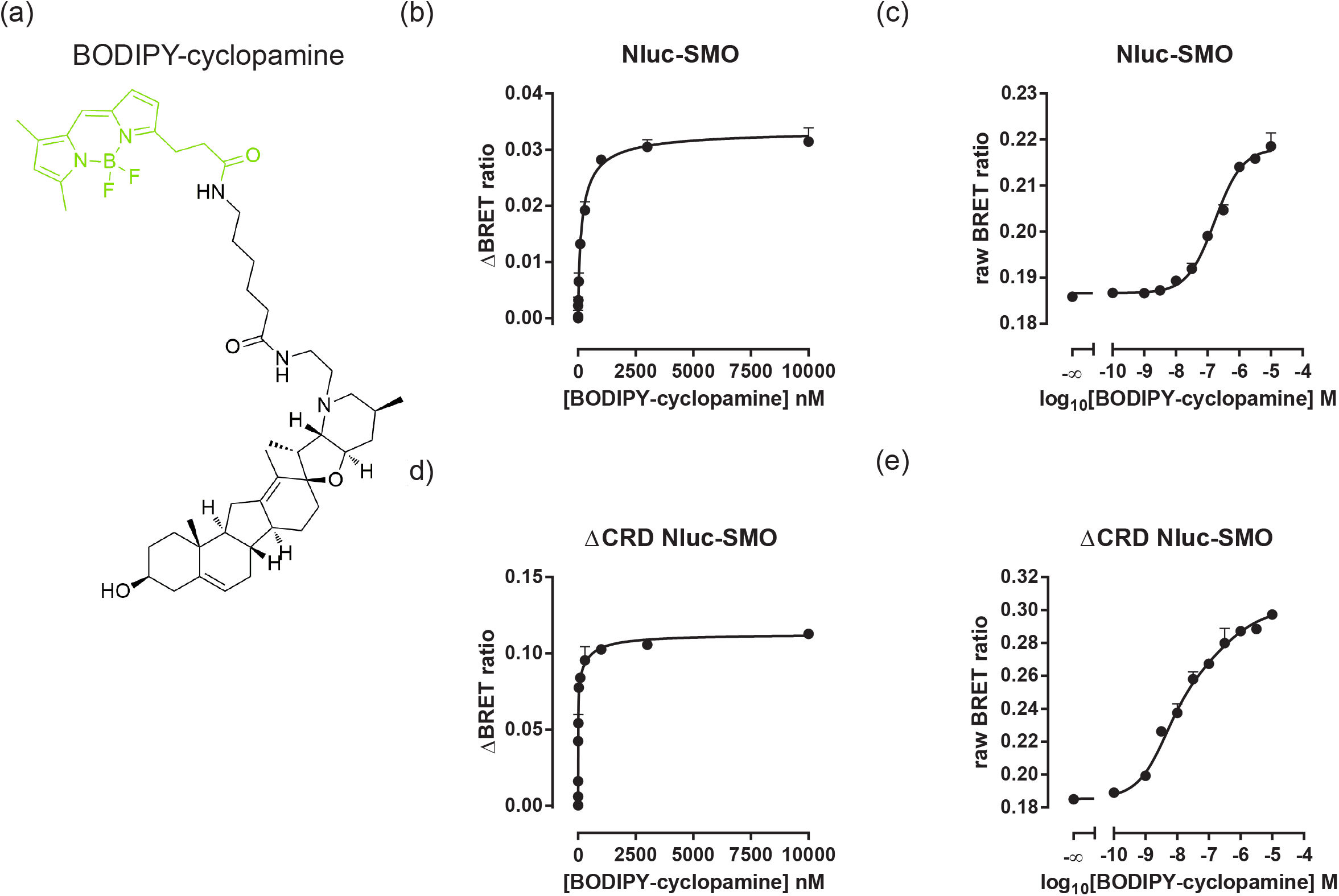
BODIPY-cyclopamine binding to SMO. (a) Chemical structure of BODIPY-cyclopamine. The BODIPY moiety is highlighted in green. The structure was drawn using ACD/ChemSketch freeware. NanoBRET BODIPY-cyclopamine assays were performed in ΔSMO HEK293 cells transiently expressing Nluc-tagged SMO (Nluc-SMO; b, c) or ΔCRD Nluc-SMO (d, e). Saturation curves are presented as hyperbolic curves with linear (b, d) and as sigmoidal curves with logarithmic (c, e) BODIPY-cyclopamine concentrations. Graphs present raw NanoBRET values obtained following 1 h ligand exposure to living ΔSMO HEK293 cells. N=8-9. Data are presented as mean±SEM. Curves for Nluc-SMO were fitted to a three parameter model (log scale) and one-site specific binding (linear scale). For ΔCRD Nluc-SMO curves were fitted according to biphasic (log scale) or two-site specific binding (linear scale).

**Fig. 3:**
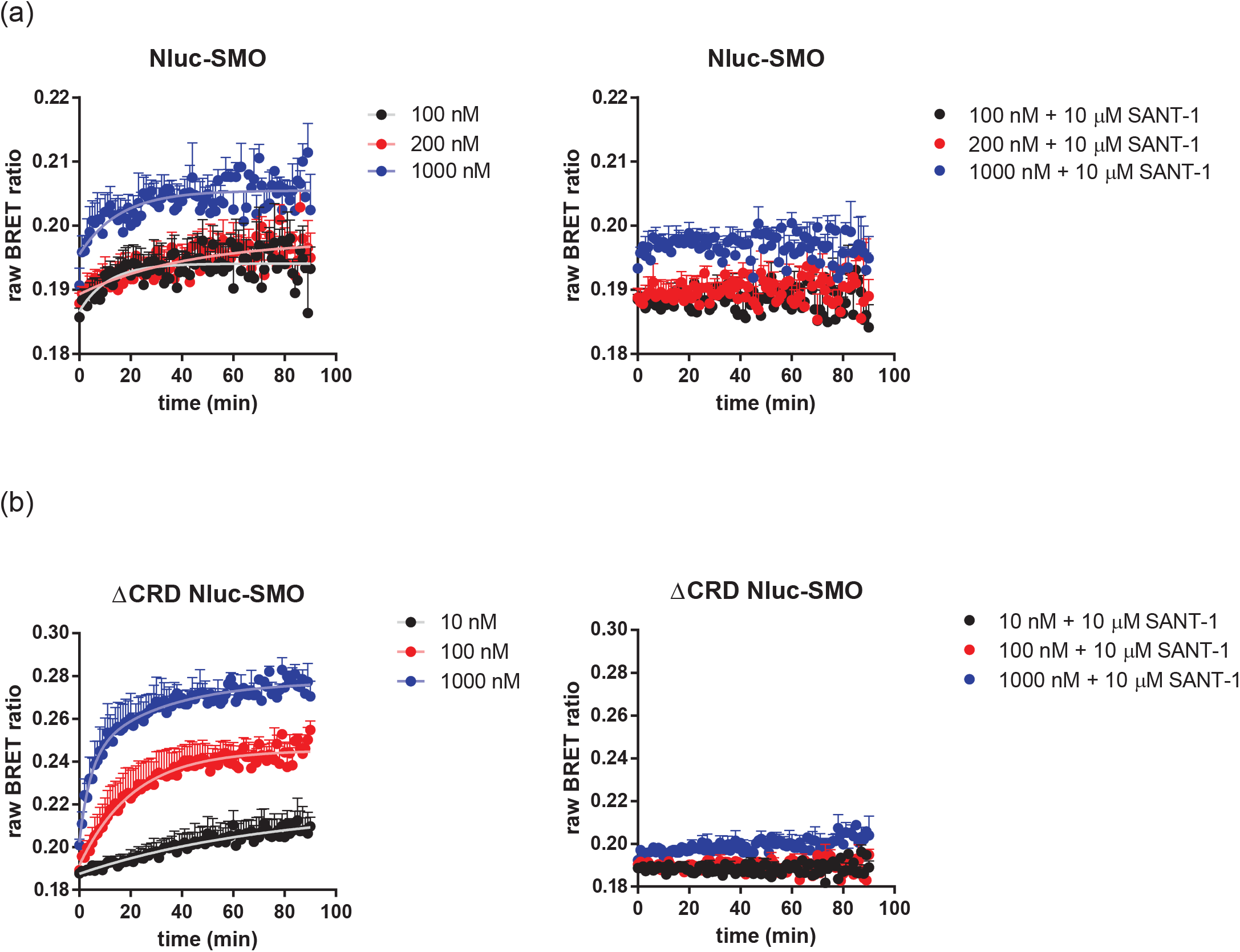
BODIPY-cylcopamine binding kinetics. Association kinetics of BODIPY-cyclopamine to Nluc-SMO (a; 100, 200 and 1000 nM BODIPY-cyclopamine) or ΔCRD Nluc-SMO (b; 10, 100 and 1000 nM BODIPY-cyclopamine) were determined in the absence and presence of the SMO antagonist SANT-1 (10 μM) by detection of NanoBRET in living ΔSMO HEK293 cells over time. NanoBRET was sampled once per minute for 90 min. Data are presented as mean±SEM from three independent experiments. Kinetic parameters are summarized in Table 3 and 4.

**Table 1:**
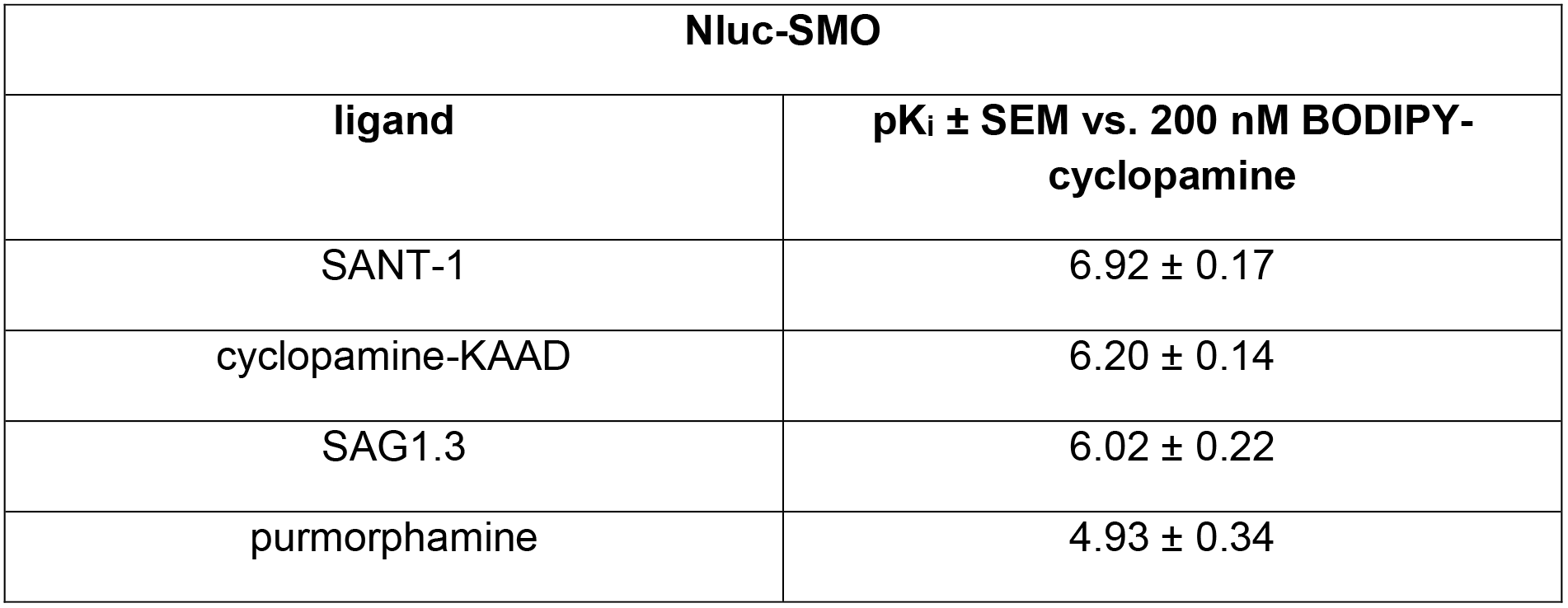
Binding affinities of various SMO ligands in competition with BODIPY-cyclopamine binding (200 nM) to Nluc-SMO. Data are based on five independent experiments presented in Fig. 5a. pK_i_ values are presented as mean ± SEM.

**Table 2:** Binding affinities of various SMO ligands in competition with BODIPY-cyclopamine binding (10 nM) to ΔCRD Nluc-SMO. Data are based on five to six independent experiments presented in Fig. 5b. pK_i_ values are presented as mean ± SEM.

| **ΔCRD Nluc-SMO** |  |
| --- | --- |
| **ligand** | **pK<sub>i</sub> ± SEM vs. 10 nM BODIPY-cyclopamine** |
| SANT-1 | 8.76 ± 0.14 |
| cyclopamine-KAAD | 8.55 ± 0.21 |
| SAG1.3 | 7.95 ± 0.15 |
| purmorphamine | 6.04 ± 0.42 |

**Table 3:** Kinetic parameters of BODIPY-cyclopamine binding to Nluc-SMO. Values are based on three independent experiments (shown in Fig. 3a).

| **Nluc-SMO** |  |  |  |  |  |  |
| --- | --- | --- | --- | --- | --- | --- |
|  | **k<sub>obs</sub> ± SEM**<br>**($\frac{1}{M \cdot min}$)** | **t<sub>1/2</sub> ± SEM**<br>**(min)** | **k<sub>obs</sub> fast ± SEM**<br>**($\frac{1}{M \cdot min}$)** | **k<sub>obs</sub> slow ± SEM**<br>**($\frac{1}{M \cdot min}$)** | **t<sub>1/2</sub> fast ± SEM**<br>**(min)** | **t<sub>1/2</sub> slow ± SEM**<br>**(min)** |
| One-phase association<br>(100 nM) | 0.0906 ± 0.0360 | 7.65 |  |  |  |  |
| One-phase association<br>(200 nM) | 0.0237 ± 0.0112 | 29.25 |  |  |  |  |
| One-phase association<br>(1000 nM) | 0.0641 ± 0.0172 | 10.82 |  |  |  |  |

**Table 4:** Kinetic parameters of BODIPY-cyclopamine binding to ΔCRD Nluc-SMO. Values are based on three independent experiments (shown in Fig. 3b).

| **ΔCRD Nluc-SMO** |  |  |  |  |  |  |
| --- | --- | --- | --- | --- | --- | --- |
|  | **k<sub>obs</sub> ± SEM**<br>**($\frac{1}{M \cdot min}$)** | **t<sub>1/2</sub> ± SEM**<br>**(min)** | **k<sub>obs</sub> fast ± SEM**<br>**($\frac{1}{M \cdot min}$)** | **k<sub>obs</sub> slow ± SEM**<br>**($\frac{1}{M \cdot min}$)** | **t<sub>1/2</sub> fast ± SEM**<br>**(min)** | **t<sub>1/2</sub> slow ± SEM**<br>**(min)** |
| One-phase association<br>(10 nM) | 0.0133 ± 0.0077 | 52.26 |  |  |  |  |
| One-phase association<br>(100 nM) | 0.0474 ± 0.0067 | 14.62 |  |  |  |  |
| Two-phase association<br>(1000 nM) |  |  | 0.2299 ± 0.1068 | 0.034 ± 0.0171 | 3.0 | 20.6 |

### NanoBRET-based ligand binding is superior to fluorescence-based quantification of BODIPY-cyclopamine binding to SMO

Cyclopamine is chemically similar to cholesterol rendering it cell permeable and lipophilic resulting in high unspecific binding to cells and particularly membranes. When comparing the increase in NanoBRET between BODIPY-cyclopamine and Nluc-SMO (or ΔCRD Nluc-SMO) with the increase in the fluorescence signal emerging from BODIPY-cyclopamine, specific and saturable binding can be detected by NanoBRET already in the lower nanomolar range especially with the ΔCRD Nluc-SMO construct. On the other hand, the non-saturable increase in fluorescence is detectable only at higher concentrations of BODIPY-cyclopamine. More importantly, the NanoBRET signal saturates at ligand concentrations that produce an unreliable increase in fluorescence, especially in the case of the ΔCRD Nluc-SMO. At higher BODIPY-cyclopamine concentrations, beyond those required to saturate the NanoBRET signal, a linear increase of fluorescence was detectable indicating that fluorescence includes a substantial component of unspecific ligand binding (Fig. 4a,b). This was particularly obvious when comparing the fluorescence signal at a BODIPY-concentration producing maximal binding in the absence and presence of 10 μM SANT-1 for the Nluc-SMO and the ΔCRD Nluc-SMO constructs (Fig. 4c). At this concentration the NanoBRET signal was blocked by SANT-1, whereas fluorescence was not affected.

**Fig. 4:**
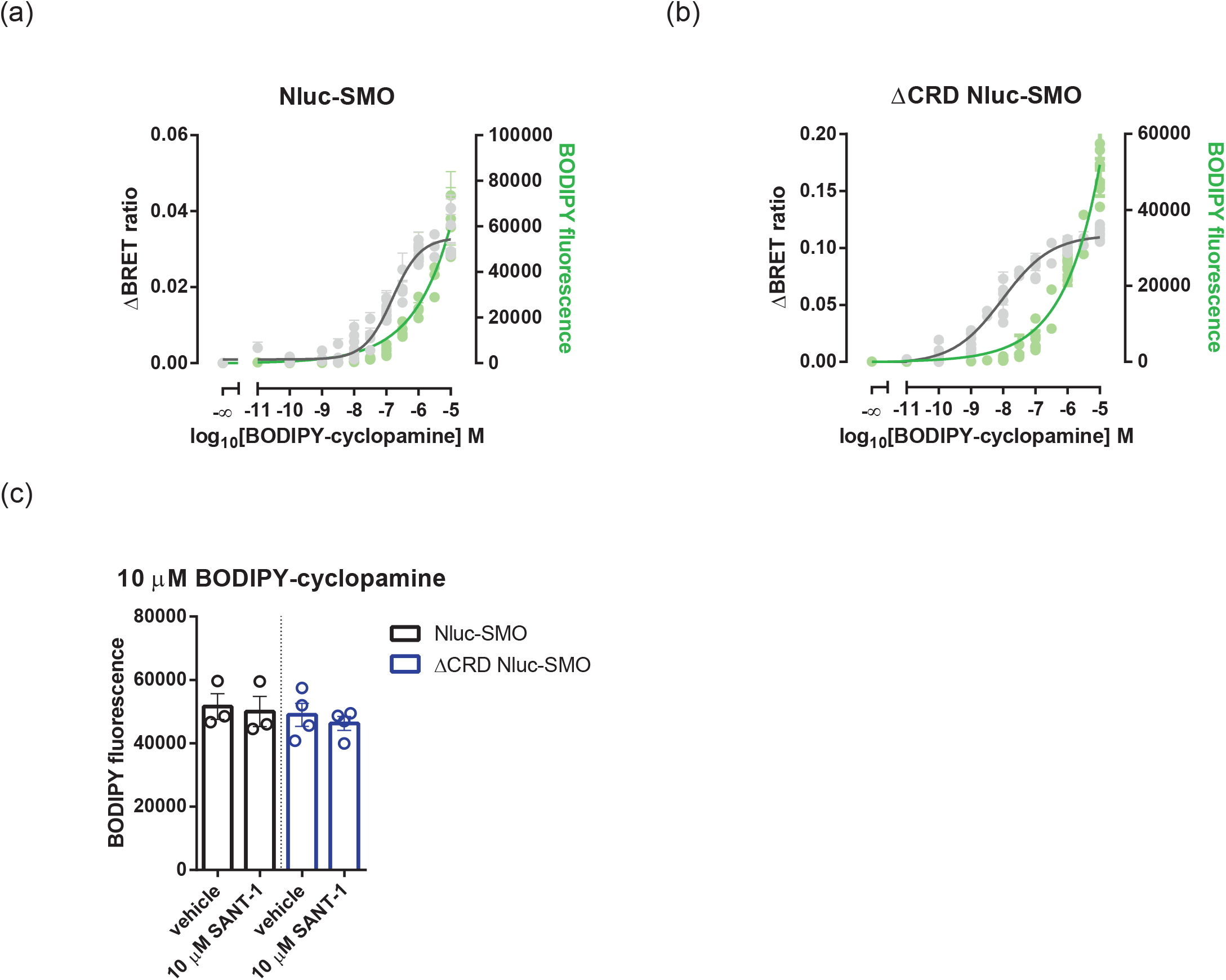
Assessment of BODIPY-cyclopamine by NanoBRET is superior to detection of fluorescence. Prior to NanoBRET binding assays with BODIPY-cyclopamine, total fluorescence values (BODIPY) were detected. The graphs present increase in NanoBRET (left axis) and BODIPY fluorescence (right axis; green) for Nluc-SMO (a) and ΔCRD Nluc-SMO (b). NanoBRET (ΔBRET) data are extracted from the experiments shown in Fig. 1. Data points are shown as mean±SD of each technical replicate of eight to nine independent experiments. Curve fitting for fluorescence values was done using semi-log line function in GraphPad Prism 6. (c) BODIPY-cyclopamine fluorescence is compared in the absence and presence of 10 μM SANT-1 in experiments with Nluc-SMO or ΔCRD Nluc-SMO. While SANT-1 dramatically affects NanoBRET readings (see Fig. 3), fluorescence values are not affected. Data present mean±SEM of three to four independent experiments.

### BODIPY-cyclopamine binding to SMO is surmountable

To explore the competitive nature of BODIPY-cyclopamine binding to SMO in more detail, we combined BODIPY-cyclopamine with increasing concentrations of commercially available SMO antagonist (SANT-1), inverse agonist (cyclopamine-KAAD) and agonists (SAG1.3 and purmorphamine) employing both the full length Nluc-SMO (competition with 200 nM BODIPY-cyclopamine; the fluorescent ligand used at this concentration gave approximately 50% of max NanoBRET in the saturation experiments) as well as the ΔCRD Nluc-SMO (competition with 10 nM BODIPY-cyclopamine; the fluorescent ligand used at this concentration gave approximately 50% of max NanoBRET in the saturation experiments). While cyclopamine-KAAD and SANT-1 presented the highest affinity to Nluc-SMO, the agonist SAG1.3 was intermediate and purmorphamine showed the lowest affinity (**Fig. 5a, Table 1**). A similar rank order was obtained in the ΔCRD Nluc-SMO-transfected cells (**Fig. 5b, Table 2**). Interestingly, residual BODIPY-cyclopamine binding produced NanoBRET, at competitor concentrations sufficiently high to reach saturation, that were substantially higher in the full length Nluc-SMO compared (**Fig. 5a**) to ΔCRD Nluc-SMO (**Fig. 5b**). At maximal competition SANT-1 reduced BODIPY-cyclopamine (200 nM) binding to 45.2% of maximal binding, whereas it completely abolished BODIPY-cyclopamine (10 nM) binding at the ΔCRD Nluc-SMO (~0.1% binding left). These findings indicate that, at the tested concentrations, BODIPY-cyclopamine binding is surmountable to a higher degree at the ΔCRD Nluc-SMO compared to Nluc-SMO. Additionally, cyclopamine-KAAD competition with BODIPY-cyclopamine at the full-length receptor did not reach the plateau indicating further displacement of the fluorescent ligand, presumably at a different binding pocket.

**Fig. 5:**
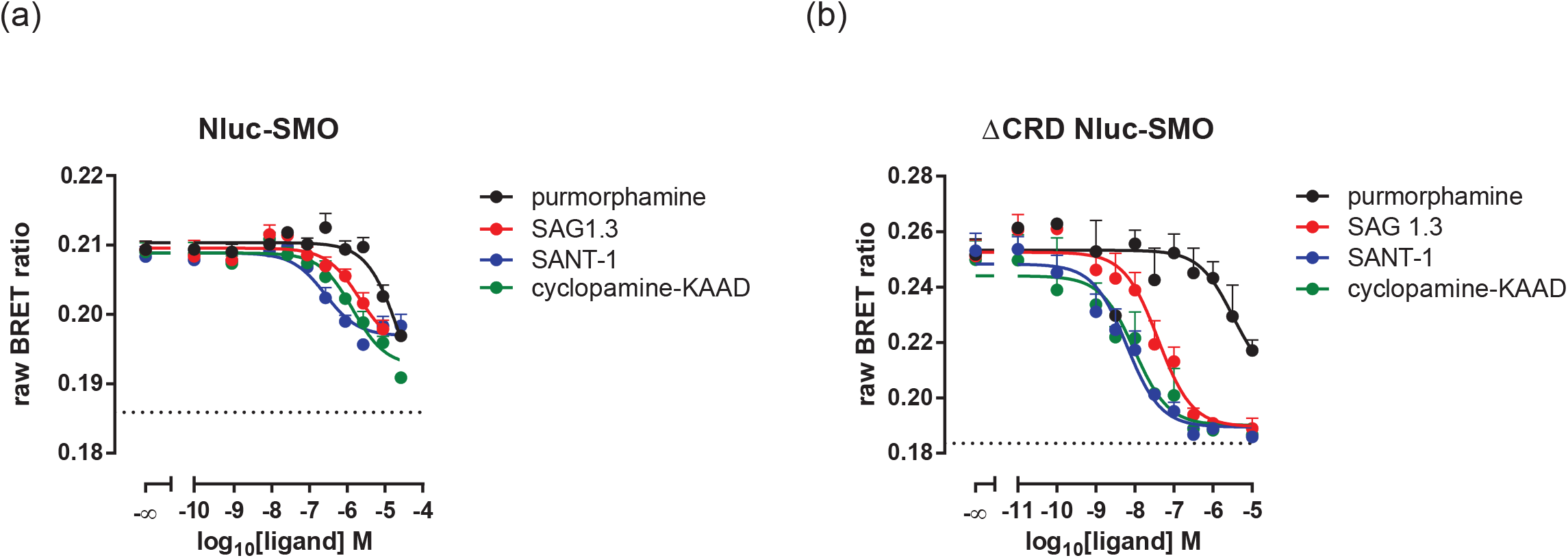
Competition experiments with BODIPY-cyclopamine and SMO antagonists and agonists. ΔSMO HEK293 cells expressing Nluc-SMO (a) or ΔCRD Nluc-SMO (b) were incubated with increasing concentrations of SMO antagonists/inverse agonists (SANT-1, cyclopamine-KAAD) and agonists (SAG1.3, purmorphamine) and subsequently exposed to BODIPY-cyclopamine (200 nM for Nluc-SMO; 10 nM for ΔCRD Nluc-SMO). Raw NanoBRET data are presented as mean±SEM from five to six independent experiments performed in technical triplicates. Pharmacological parameters are summarized in Table 1 and 2. Curve fitting was done with a one-site competition binding model. The dotted lines represent raw NanoBRET ratio of the donor-only condition (no BODIPY-cyclopamine added).

### Differential competition of BODIPY-cyclopamine binding allows pharmacological separation of two binding sites

The successful targeting of SMO with vismodegib in the therapy of basal cell carcinoma has also led to the discovery of therapy-resistant point mutations in SMO (Zhang, Sun, Liu & Song, 2018). Here, we introduced the double mutant D477G^6.54^/E522K^7.38^ into Nluc-SMO and ΔCRD Nluc-SMO (corresponding to human D473^6.54^ and E518^7.38^ mutants) in order to further dissect the contribution of different binding sites to BODIPY-cyclopamine-SMO interaction. The mutant versions are expressed on the cell surface upon transient transfection in DSMO HEK293 cells (**Fig. 6a**). In order to further define the binding characteristics of the two separate BODIPY-cyclopamine binding sites in the 7TM core of the receptor, we made use of a saturating concentration of SANT-1 (10 μM) and probed the wild type and D477G^6.54^/E522K^7.38^ of Nluc-SMO and ΔCRD Nluc-SMO with increasing concentrations of BODIPY-cyclopamine (**Fig. 6b, c**). In line with the competition data using a fixed BODIPY-cyclopamine concentration, we found that SANT-1, which solely binds to the 7TM ligand binding site of SMO (Wang et al., 2014), reduces the maximal binding of BODIPY-cyclopamine at the Nluc-SMO by one third with maintained affinity. At the ΔCRD Nluc-SMO, however, SANT-1 virtually prevents BODIPY-cyclopamine interaction with SMO up to a concentration of 100 nM. At 100 nM and above BODIPY-cyclopamine reliably showed specific, SANT-1 (10 μM)-insensitive binding in cells transfected with ΔCRD Nluc-SMO. The SANT-1-insensitive fraction of BODIPY-cyclopamine shows a reduced B_max_ and a lower affinity. In the full length Nluc-SMO, the double mutant did not affect B_max_ of BODIPY-cyclopamine and affinity was only slightly reduced (one-site fit B_max_ = 0.035 ± 0.002, K_d_ = 666.2 ± 121.8 nM). In ΔCRD Nluc-SMO D477G^6.54^/E522K^7.38^, on the other hand, the BODIPY-cyclopamine curve was clearly right-shifted and maximal binding was reduced (one-site fit B_max_ = 0.074 ± 0.001, K_d_ = 450.2 ± 38.7 nM). In both cases, SANT-1 (10 μM), which targets the lower binding pocket, maintains its effect on BODIPY-cyclopamine at both the wt and the D477G^6.54^/E522K^7.38^ for full length and ΔCRD SMO. Importantly, the double mutant does not affect the SANT-1-insensitive fraction of BODIPY-cyclopamine binding in ΔCRD Nluc-SMO (**Fig. 6c**). In addition, we provide a kinetics analysis of BODIPY-cyclopamine association to D477G^6.54^/E522K^7.38^ ΔCRD Nluc-SMO (**Fig. 6d**; **Table 5**). At 1000 nM BODIPY-cyclopamine the t_1/2_ and k_obs_ of the D477G^6.54^/E522K^7.38^ ΔCRD Nluc-SMO are very similar to Nluc-SMO, suggesting binding to the same ligand binding site.

**Fig. 6:**
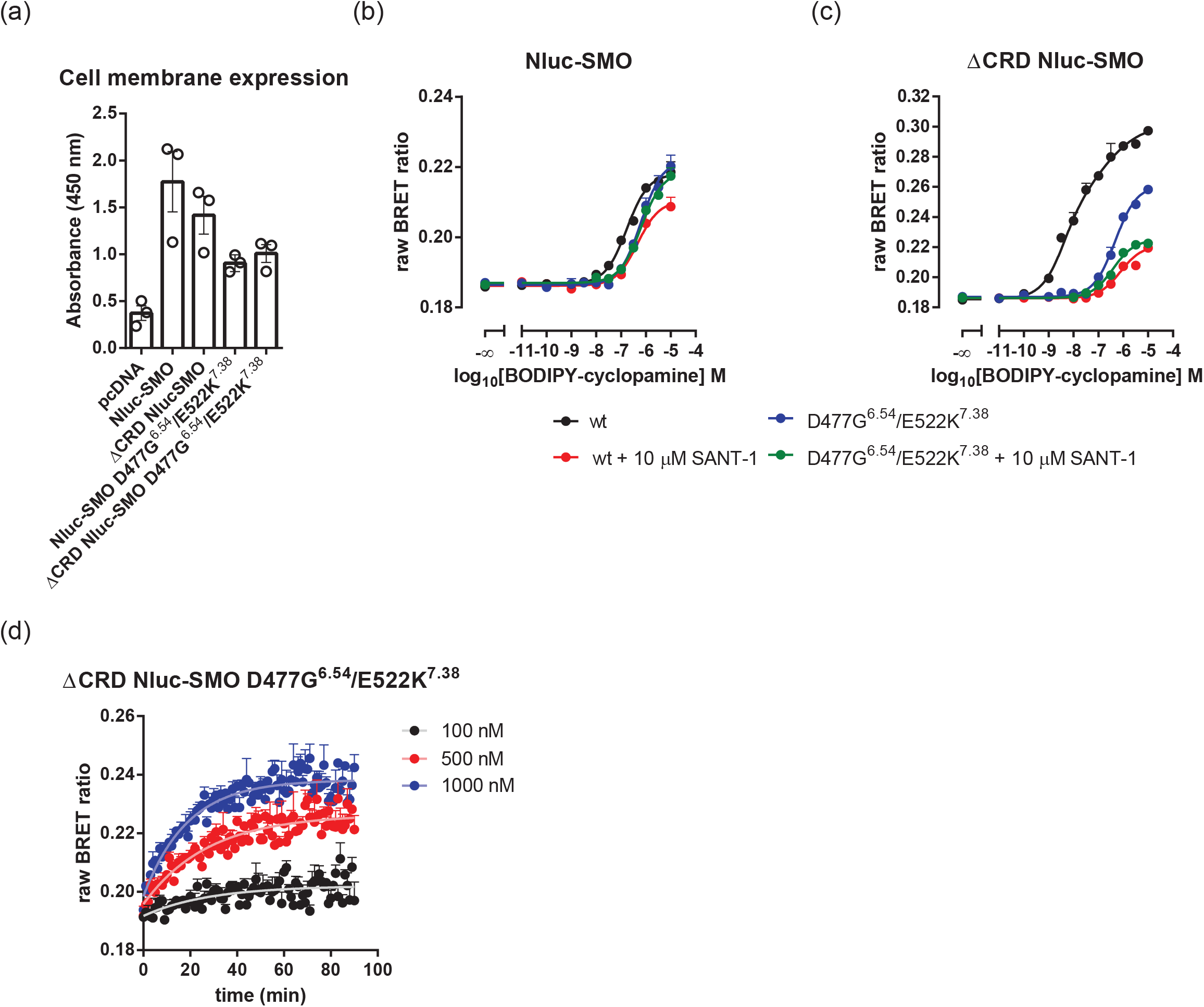
Combining BODIPY-Cyclopamine with SANT-1 allows pharmacological dissection of two separate binding sites in the 7TM core of SMO. (a) Surface ELISA was used to assess cell surface expression of wild type and D477G^6.54^/E522K^7.38^ double mutant of Nluc-SMO and ΔCRD Nluc-SMO upon overexpression in ΔSMO HEK293 cells. Data are shown as mean±SEM of three independent experiments performed in technical triplicates. BODIPY-cyclopamine binding curves were determined in ΔSMO HEK293 cells expressing either Nluc-SMO (b) or ΔCRD Nluc-SMO (c) either in the absence or presence of a saturating concentration of the SMO antagonist SANT-1 (10 μM), which targets the lower pocket of the 7TM ligand binding site of SMO. Data are presented as mean±SEM of three to five independent experiments performed in technical duplicates or triplicates. Values for the wild type SMO in the absence of SANT-1 are from Fig. 2c (Nluc-SMO) and Fig. 2e (ΔCRD Nluc-SMO). (d) Association kinetics of BODIPY-cyclopamine association to ΔCRD Nluc-SMO D477G^6.54^/E522K^7.38^. Data from three independent experiments performed in technical triplicates are presented as mean±SEM. Kinetic parameters are summarized in Table 5.

**Table 5:** Kinetic parameters of BODIPY-cyclopamine binding to ΔCRD Nluc-SMO D477G^6.54^/E522K^7.38^. Values are based on three independent experiments (shown in Fig. 6d).

| **ΔCRD Nluc-SMO D477G<sup>6.54</sup>/E522K<sup>7.38</sup>** |  |  |  |  |  |  |
| --- | --- | --- | --- | --- | --- | --- |
|  | **$k_{obs} \pm$**<br>**SEM**<br>**$(\frac{1}{M*min})$** | **$t_{1/2} \pm$**<br>**SEM**<br>**(min)** | **$k_{obs}$**<br>**fast ±**<br>**SEM**<br>**$(\frac{1}{M*min})$** | **$k_{obs}$ slow**<br>**± SEM**<br>**$(\frac{1}{M*min})$** | **$t_{1/2}$ fast**<br>**± SEM**<br>**(min)** | **$t_{1/2}$**<br>**slow**<br>**± SEM**<br>**(min)** |
| One-phase<br>association<br>(100 nM) | 0.0356 ±<br>0.0121 | 19.44 | | | | |
| One-phase<br>association<br>(500 nM) | 0.0363<br>± 0.0046 | 19.09 | | | | |
| One-phase<br>association<br>(1000 nM) | 0.0536<br>±<br>0.0046 | 12.94 | | | | |

## 4. DISCUSSION

The development of a NanoBRET-based ligand binding assay provides an interesting complement to GPCR pharmacology enabling ligand binding studies on living cells in real time with simplified protocols (Stoddart et al., 2015; Stoddart, Kilpatrick & Hill, 2018). Here, we optimize this assay for the Class F receptor SMO improving sensitivity and performance of previously used fluorescence-based approaches (Chen, Taipale, Cooper & Beachy, 2002; Chen, Taipale, Young, Maiti & Beachy, 2002; Gorojankina et al., 2013; Huang et al., 2016; Huang et al., 2018; Manetti et al., 2010; Tao et al., 2011). Due to its large assay window for specific binding and the low influence of unspecific binding this NanoBRET-based assay is particularly suitable for lipophilic ligands such as cyclopamine and potentially other cholesterol-like molecules, which target SMO and generally show unspecific interactions with the membrane. Furthermore, this high-throughput compatible assay should be adaptable to any fluorescently-tagged molecule acting as SMO ligand and could – provided small molecules become available to target Frizzleds also be employed for other Class F receptors.

Recent insight into the molecular mechanisms of drug action on SMO by crystallography and CryoEM provide somewhat controversial yet intriguing information regarding cyclopamine and cholesterol interaction with the 7TM ligand-binding site and the CRD (Deshpande et al., 2019; Huang et al., 2016; Huang et al., 2018; Qi, Liu, Thompson, McDonald, Zhang & Li, 2019; Weierstall et al., 2014). Here, we have been able to pharmacologically separate two BODIPY-cyclopamine binding sites on the 7TM core of SMO. Total BODIPY-cyclopamine binding to full length Nluc-SMO is composed of at least two components: ligand binding to the CRD (Huang et al., 2016; Huang et al., 2018) – for which we did not find evidence for in our experiments – and the 7TM core (Huang et al., 2018; Weierstall et al., 2014). Most importantly, SANT-1, which solely binds to the receptor core in the lower pocket of the SMO binding site (Wang et al., 2014), competes with BODIPY-cyclopamine more efficiently in the ΔCRD Nluc-SMO compared to the full length receptor. This large increase in affinity and NanoBRET signal/B_max_ suggest that the CRD exerts a negative allosteric modulation on BODIPY-cyclopamine binding to the SMO 7TM core. The residual binding of BODIPY-cyclopamine to ΔCRD Nluc-SMO above 10^−7^ M identifies a SANT-1-insensitive fraction suggesting an additional binding pocket for BODIPY-cyclopamine. Given the simultaneous binding of SAG21k and cholesterol to SMO in the recent crystal structure (PDB 6O3C) of active, nanobody NbSmo8-bound SMO (Deshpande et al., 2019), and the fact that SANT-1 and cyclopamine occupy two different parts of the small molecule binding space in SMO (Huang et al., 2018; Wang et al., 2014; Weierstall et al., 2014) it could also be possible that BODIPY-cyclopamine and SANT-1 bind the 7TM course simultaneously. It remains to be verified by structural studies if binding modes of cyclopamine and BODIPY-cyclopamine are indeed the same. Along these lines, different binding poses of cyclopamine and related sterols were reported in several crystal structures using SMO from various species. In summary, there is a cyclopamine/sterol binding site in the lipid groove of the CRD (Byrne et al., 2016; Deshpande et al., 2019; Huang et al., 2016; Huang et al., 2018), one in the upper pocket of the 7TM core (Huang et al., 2018; Qi, Liu, Thompson, McDonald, Zhang & Li, 2019) and one in the lower position (Deshpande et al., 2019), which overlaps with the SANT-1 binding pocket (Wang et al., 2014) (**Fig. 7**). Moreover, SAG1.3-induced Gli transcriptional activity still reaches saturation albeit at lower efficacy following treatment with SANT-1.This further provides functional evidence that both the upper (SAG1.3 binding site) and the lower pocket (SANT-1 binding site) in the 7TM core of SMO can be occupied by ligands simultaneously and that these pockets may allosterically regulate each other (Chen, Taipale, Young, Maiti & Beachy, 2002). Similar conclusions were drawn from radioligand binding studies (Frank-Kamenetsky et al., 2002; Tao et al., 2011). Furthermore, the phenomenon of allosteric regulation in the SMO binding site was also inferred from studies on other SMO ligands (Chen et al., 2016; Hoch et al., 2015).

**Fig. 7:**
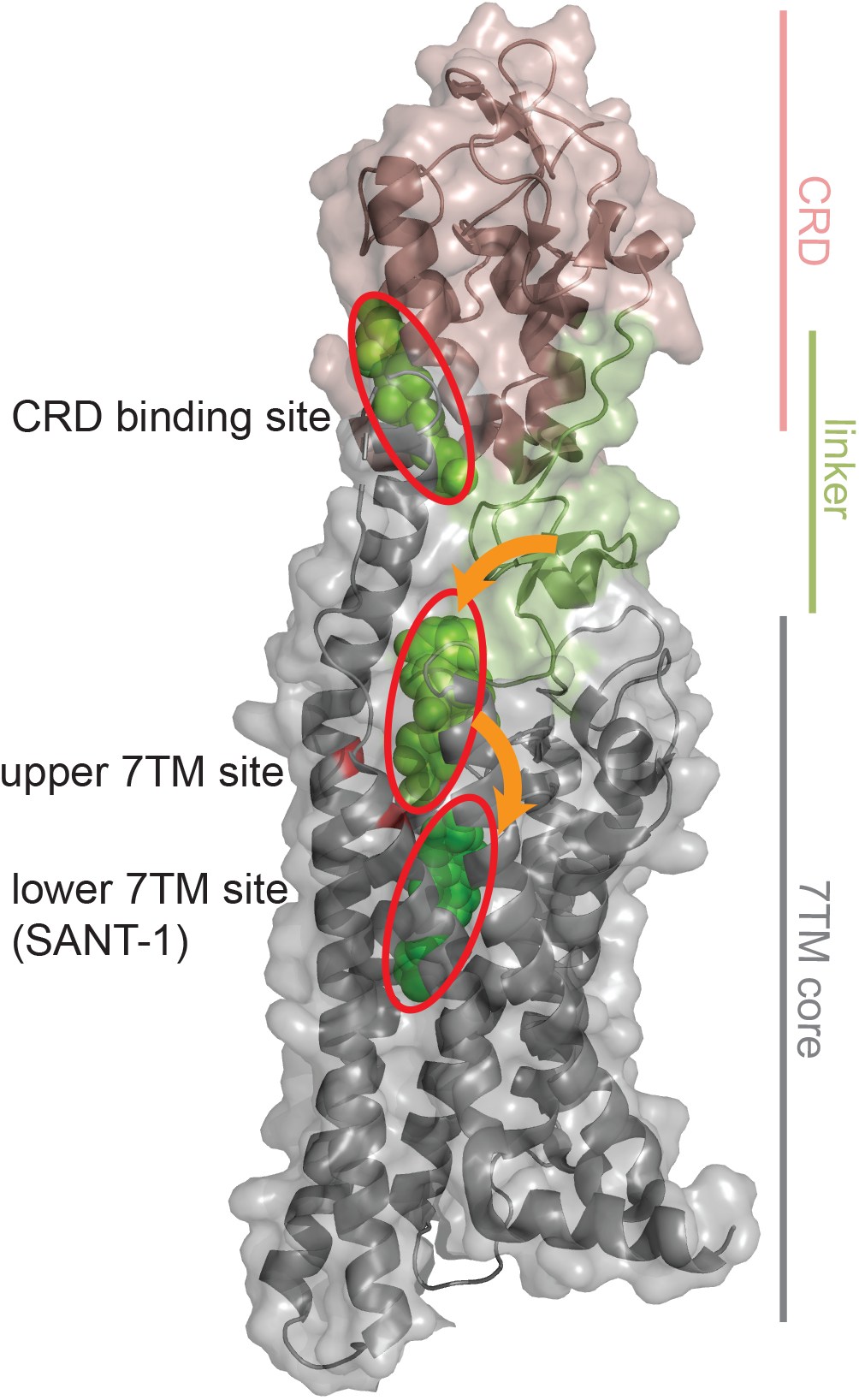
Separate ligand binding sites on SMO. Schematic presentation of reported SMO ligand binding sites. The receptor model (transparent surface view with protein backbone shown as ribbon) was derived from the SMO crystal structure bound to Nanobody NbSmo8 and SAG21k (PDB ID: 6O3C) (Deshpande et al., 2019). The major subdomains (CRD, linker and 7TM core) are color-coded. Ligands are shown as spheres and are highlighted in green. Reported ligand binding sites on the CRD and in the 7TM core are encircled in red. While our data did not indicate BODIPY-cyclopamine binding to the CRD site, the NanoBRET-based binding assay supports two communicating binding sites for BODIPY-cyclopamine in the 7TM core of SMO. The mutations D477G^6.54^/E522K^7.38^ in TM6 and TM7, which affect BODIPY-cyclopamine binding to SMO are depicted in red. Based on the effects of D477G^6.54^/E522K^7.38^ and SANT-1 on BODIPY-cyclopamine binding we suggest a two step binding mode of BODIPY-cyclopamine involving the upper and the lower 7TM site (orange arrows). The SMO model was produced in PyMOL (The PyMOL Molecular Graphics System, Version 2.0 Schrödinger, LLC).

Interestingly, removal of the CRD of SMO increased both B_max_ and the K_d_ of BODIPY-cyclopamine. The absence of the CRD, which obviously includes extinction of the proposed CRD binding site for BODIPY-cyclopamine, most likely provides better access to the normally buried binding site in the core of the receptor causing a left shift and an increased B_max_. Related to that, one might also postulate an allosteric action of the CRD on the ligand-binding site on the 7TM core, an idea that is fueled by the observation that ΔCRD SMO exerts a higher constitutive activity (Byrne et al., 2016). A part of the explanation for the efficacy shift of the BODIPY-cyclopamine binding curve could also be the altered BRET parameters since the distance of the NanoBRET donor and the acceptor are likely shorter in the ΔCRD SMO compared to the full length receptor. However, while the different BRET efficiencies in the two receptor constructs could affect the amplitude of the NanoBRET signal originating from BODIPY-cyclopamine binding, it cannot explain the leftward shift of the binding curves indicating a higher affinity to the ΔCRD Nluc-SMO. It should be noted that in previous publications low concentrations of BODIPY-cyclopamine (usually 5 nM) were used for SMO binding assays. Given our data regarding the different affinities of the separate BODIPY-cyclopamine binding sites, these low ligand concentrations neither allowed sampling the SANT-1-insensitive, low affinity site in the core of ΔCRD-Nluc SMO nor the putative site on the CRD of the full-length receptor. Application of the novel NanoBRET methodology, however, allows to reliably detect even picomolar and nanomolar amounts of BODIPY-cyclopamine bound to ΔCRD Nluc-SMO and ΔCRD Nluc-SMO D477G^6.54^/E522K^7.38^.

In summary, we dissect BODIPY-cyclopamine interaction with two binding sites on the 7TM core with different affinities. The high affinity site is the deep SANT-1 binding pocket. Since the D477G^6.54^/E522K^7.38^ in TM6 and TM7 partially affect the high affinity component of BODIPY-cyclopamine binding, it could be that interaction with D477^6.54^/E522^7.38^ provides a transition mechanism of ligand binding to the deeper pocket. Low affinity binding to the D477G^6.54^/E522K^7.38^ mutant in the upper 7TM site is still possible and that binding is insensitive to allosteric modulation by SANT-1 binding to the deeper site.

For drug discovery efforts, the ΔCRD Nluc-SMO probe presents a valuable tool with an advantageous assay window. The intrinsic caveat of the lack of the physiologically relevant CRD, however, remains, requiring thorough validation of screening hits in assays relying on full length SMO. Further work will address in which way the separate ligand-binding sites interact allosterically and what role the CRD-core contacts play for that potential communication.

## ACKNOWLEDGEMENTS

We thank Stephen Hill (University of Nottingham, UK) for the Nluc-A3 construct, Anna Krook (Karolinska Institutet, Sweden) for the access to the ClarioStar plate reader and Ainoleena Turku (Karolinska Institutet, Sweden) for drawing the chemical structure of BODIPY-cyclopamine. The study was supported by grants from Karolinska Institutet, the Swedish Research Council (2017-04676), the Swedish Cancer Society (CAN2017/561), the Novo Nordisk Foundation (NNF17OC0026940), Stiftelsen Olle Engkvist Byggmästare (2016/193), Wenner Gren Foundations (UPD2018-0064), Emil and Wera Cornells Stiftelse.

## CONFLICT OF INTEREST

The authors declare no conflicts of interest.

## AUTHOR CONTRIBUTIONS

P.K. and G.S. designed the research. P.K. and C.-F.B. generated the ΔCRD and full-length Nluc-SMO constructs, conducted experiments and analyzed the data. P.K., C.-F.B. and G.S. wrote or contributed to the writing of the manuscript.

## References

Ballesteros JA, & Weinstein H (1995). Integrated methods for the construction of three-dimensional models and computational probing of structure-function relations in G protein-coupled receptors. Methods in Neurosciences 25: 366–428.

Bee WT, Xie W, Truong M, Will M, Turunen B, Zuercher WJ, et al. (2012). The development of a high-content screening binding assay for the smoothened receptor. J Biomol Screen 17: 900–911.

Bosma R, Stoddart LA, Georgi V, Bouzo-Lorenzo M, Bushby N, Inkoom L, et al. (2019). Probe dependency in the determination of ligand binding kinetics at a prototypical G protein-coupled receptor. Scientific reports 9: 7906.

Bouzo-Lorenzo M, Stoddart LA, Xia L, Ap IJ, Heitman LH, Briddon SJ, et al. (2019). A live cell NanoBRET binding assay allows the study of ligand-binding kinetics to the adenosine A3 receptor. Purinergic Signal.

Byrne EF, Sircar R, Miller PS, Hedger G, Luchetti G, Nachtergaele S, et al. (2016). Structural basis of Smoothened regulation by its extracellular domains. Nature 535: 517–522.

Chen B, Trang V, Lee A, Williams NS, Wilson AN, Epstein EH, Jr., et al. (2016). Posaconazole, a Second-Generation Triazole Antifungal Drug, Inhibits the Hedgehog Signaling Pathway and Progression of Basal Cell Carcinoma. Mol Cancer Ther 15: 866–876.

Chen JK (2016). I only have eye for ewe: the discovery of cyclopamine and development of Hedgehog pathway-targeting drugs. Nat Prod Rep 33: 595–601.

Chen JK, Taipale J, Cooper MK, & Beachy PA (2002). Inhibition of Hedgehog signaling by direct binding of cyclopamine to Smoothened. Genes Dev 16: 2743–2748.

Chen JK, Taipale J, Young KE, Maiti T, & Beachy PA (2002). Small molecule modulation of Smoothened activity. Proc Natl Acad Sci U S A 99: 14071–14076.

Cheng Y, & Prusoff WH (1973). Relationship between the inhibition constant (K1) and the concentration of inhibitor which causes 50 per cent inhibition (I50) of an enzymatic reaction. Biochem Pharmacol 22: 3099–3108.

Deshpande I, Liang J, Hedeen D, Roberts KJ, Zhang Y, Ha B, et al. (2019). Smoothened stimulation by membrane sterols drives Hedgehog pathway activity. Nature.

Frank-Kamenetsky M, Zhang XM, Bottega S, Guicherit O, Wichterle H, Dudek H, et al. (2002). Small-molecule modulators of Hedgehog signaling: identification and characterization of Smoothened agonists and antagonists. J Biol 1: 10.

Gorojankina T, Hoch L, Faure H, Roudaut H, Traiffort E, Schoenfelder A, et al. (2013). Discovery, molecular and pharmacological characterization of GSA-10, a novel small-molecule positive modulator of Smoothened. Mol Pharmacol 83: 1020–1029.

Hoch L, Faure H, Roudaut H, Schoenfelder A, Mann A, Girard N, et al. (2015). MRT-92 inhibits Hedgehog signaling by blocking overlapping binding sites in the transmembrane domain of the Smoothened receptor. FASEB J 29: 1817–1829.

Huang P, Nedelcu D, Watanabe M, Jao C, Kim Y, Liu J, et al. (2016). Cellular Cholesterol Directly Activates Smoothened in Hedgehog Signaling. Cell 166: 1176–1187 e1114.

Huang P, Zheng S, Wierbowski BM, Kim Y, Nedelcu D, Aravena L, et al. (2018). Structural Basis of Smoothened Activation in Hedgehog Signaling. Cell 174: 312–324 e316.

Incardona JP, Gaffield W, Kapur RP, & Roelink H (1998). The teratogenic Veratrum alkaloid cyclopamine inhibits sonic hedgehog signal transduction. Development 125: 3553–3562.

Ingham PW, & McMahon AP (2001). Hedgehog signaling in animal development: paradigms and principles. Genes Dev 15: 3059–3087.

Manetti F, Faure H, Roudaut H, Gorojankina T, Traiffort E, Schoenfelder A, et al. (2010). Virtual screening-based discovery and mechanistic characterization of the acylthiourea MRT-10 family as smoothened antagonists. Mol Pharmacol 78: 658–665.

Mocking TAM, Verweij EWE, Vischer HF, & Leurs R (2018). Homogeneous, Real-Time NanoBRET Binding Assays for the Histamine H3 and H4 Receptors on Living Cells. Mol Pharmacol 94: 1371–1381.

Nachtergaele S, Mydock LK, Krishnan K, Rammohan J, Schlesinger PH, Covey DF, et al. (2012). Oxysterols are allosteric activators of the oncoprotein Smoothened. Nat Chem Biol 8: 211–220.

Qi X, Liu H, Thompson B, McDonald J, Zhang C, & Li X (2019). Cryo-EM structure of oxysterol-bound human Smoothened coupled to a heterotrimeric Gi. Nature.

Raleigh DR, Sever N, Choksi PK, Sigg MA, Hines KM, Thompson BM, et al. (2018). Cilia-Associated Oxysterols Activate Smoothened. Mol Cell 72: 316–327 e315.

Rominger CM, Bee WL, Copeland RA, Davenport EA, Gilmartin A, Gontarek R, et al. (2009). Evidence for allosteric interactions of antagonist binding to the smoothened receptor. J Pharmacol Exp Ther 329: 995–1005.

Schulte G (2010). International Union of Basic and Clinical Pharmacology. LXXX. The class Frizzled receptors. Pharmacol Rev 62: 632–667.

Schulte G, & Kozielewicz P (2019). Structural insight into Class F receptors - what have we learnt regarding agonist-induced activation? Basic Clin Pharmacol Toxicol.

Sever N, Mann RK, Xu L, Snell WJ, Hernandez-Lara CI, Porter NA, et al. (2016). Endogenous B-ring oxysterols inhibit the Hedgehog component Smoothened in a manner distinct from cyclopamine or side-chain oxysterols. Proc Natl Acad Sci U S A 113.

Stoddart LA, Johnstone EK, Wheal AJ, Goulding J, Robers MB, Machleidt T, et al. (2015). Application of BRET to monitor ligand binding to GPCRs. Nat Methods 12: 661–663.

Stoddart LA, Kilpatrick LE, & Hill SJ (2018). NanoBRET Approaches to Study Ligand Binding to GPCRs and RTKs. Trends Pharmacol Sci 39: 136–147.

Stoddart LA, Vernall AJ, Bouzo-Lorenzo M, Bosma R, Kooistra AJ, de Graaf C, et al. (2018). Development of novel fluorescent histamine H1-receptor antagonists to study ligand-binding kinetics in living cells. Scientific reports 8: 1572.

Sykes DA, Stoddart LA, Kilpatrick LE, & Hill SJ (2019). Binding kinetics of ligands acting at GPCRs. Mol Cell Endocrinol 485: 9–19.

Taipale J, Chen JK, Cooper MK, Wang BL, Mann RK, Milenkovic L, et al. (2000). Effects of oncogenic mutations in Smoothened and Patched can be reversed by cyclopamine. Nature 406: 1005–1009.

Tao H, Jin Q, Koo DI, Liao X, Englund NP, Wang Y, et al. (2011). Small molecule antagonists in distinct binding modes inhibit drug-resistant mutant of smoothened. Chemistry & biology 18: 432–437.

Wang C, Wu H, Evron T, Vardy E, Han GW, Huang XP, et al. (2014). Structural basis for Smoothened receptor modulation and chemoresistance to anticancer drugs. Nature communications 5: 4355.

Weierstall U, James D, Wang C, White TA, Wang D, Liu W, et al. (2014). Lipidic cubic phase injector facilitates membrane protein serial femtosecond crystallography. Nature communications 5: 3309.

Wright SC, Kozielewicz P, Kowalski-Jahn M, Petersen J, Bowin CF, Slodkowicz G, et al. (2019). A conserved molecular switch in Class F receptors regulates receptor activation and pathway selection. Nature communications 10: 667.

Wu F, Zhang Y, Sun B, McMahon AP, & Wang Y (2017). Hedgehog Signaling: From Basic Biology to Cancer Therapy. Cell Chem Biol 24: 252–280.

Zhang H, Sun Z, Liu Z, & Song C (2018). Overcoming the emerging drug resistance of smoothened: an overview of small-molecule SMO antagonists with antiresistance activity. Future Med Chem 10: 2855–2875.

Zhang Y, Bulkley DP, Xin Y, Roberts KJ, Asarnow DE, Sharma A, et al. (2018). Structural Basis for Cholesterol Transport-like Activity of the Hedgehog Receptor Patched. Cell 175: 1352–1364 e1314.

